# The effect of deep brain stimulation on cortico-subcortical networks in Parkinson’s disease patients with freezing of gait: Exhaustive exploration of a basic model

**DOI:** 10.1101/2023.02.27.529710

**Authors:** Mariia Popova, Arnaud Messé, Alessandro Gulberti, Christian Gerloff, Monika Pötter-Nerger, Claus C Hilgetag

## Abstract

Current treatments of Parkinson’s disease (PD) have limited efficacy in alleviating freezing of gait (FoG). In this context, concomitant deep brain stimulation (DBS) of the subthalamic nucleus (STN) and the substantia nigra pars reticulata (SNr) has been suggested as a potential therapeutic approach. However, the mechanisms underlying this approach are unknown. While the current rationale relies on network-based hypotheses of intensified disinhibition of brainstem locomotor areas to facilitate the release of gait motor programs, it is still unclear how simultaneous high-frequency DBS in two interconnected basal ganglia nuclei affects large-scale cortico-subcortical network activity. Here, we use a basic model of neural excitation, the susceptible-excited-refractory (SER) model, to compare effects of different stimulation modes of the network underlying FoG. We develop a network-based computational framework to compare subcortical DBS targets through exhaustive analysis of the brain attractor dynamics in the healthy, PD and DBS states. We demonstrate the validity of the approach and the superior performance of combined STN+SNr DBS in the normalization of spike propagation flow in the FoG network. The framework aims to move towards a mechanistic understanding of the network effects of DBS and may be applicable to further perturbation-based therapies of brain disorders.

**AUTHOR SUMMARY:** Parkinson’s disease patients with Freezing of Gait (FoG) may be treated by deep brain stimulation, which produces effects mediated by brain networks. Currently, the approach of combined DBS of the subthalamic nucleus and the substantia nigra pars reticulata is investigated for being particularly beneficial for patients with axial symptoms, but the exact mechanisms of this effect are unknown. Here, we present a network-based computational framework using a basic excitable model that enables us to simulate the complete activity patterns of the brain network involved in FoG. These simulations reveal network mechanisms underlying STN+SNr DBS and its efficacy in alleviating FoG. The proposed framework can capture the influence of the DBS target sites on cortico-subcortical networks and help to identify suitable stimulation targets.

## INTRODUCTION

Parkinson’s disease is a progressive neurodegenerative disorder with cardinal motor symptoms of axial and limb bradykinesia, rest tremor, and rigidity (Postuma et al., 2015). Many PD symptoms are successfully treated pharmacologically (Fox et al., 2018) or by **deep brain stimulation** (Deuschl et al., 2022; Deuschl, Paschen, & Witt, 2013). However, axial symptoms, such as parkinsonian gait disorder, postural instability and freezing of gait show limited response to treatment (Pötter-Nerger & Volkmann, 2013; Schlenstedt et al., 2017). FoG is associated with an increased risk of falls among patients and represents a significant source of morbidity. Thus, there is a need to reconcile FoG network pathophysiology and efficient treatment approaches.

DBS is an established treatment strategy for PD patients with motor fluctuations, medically refractory tremor or averse dopaminergic drug reactions (Pollak, 2013). Despite widespread use of DBS in the clinical routine, the mechanism of its action remains poorly understood. DBS efficacy varies widely among patients, and particularly patients with gait disorders and FoG appear to respond better to specific stimulation patterns (Pötter-Nerger & Volkmann, 2013). It was recently shown that simultaneous deep brain stimulation of STN and SNr (STN+SNr DBS) outperforms standard STN DBS in improving FoG symptoms (M. A. Horn et al., 2022; Wagner et al., 2022; Weiss et al., 2013). However, there is a possibility of worsening akinesia (Weiss et al., 2013), hypomania (Ulla et al., 2011), mania (Ulla et al., 2006), and depression (Blomstedt et al., 2008) with SNr stimulation, partly due to projections of the SNr to the limbic system (Pötter-Nerger & Volkmann, 2013). Therefore, while it is an attractive alternative to induce co-stimulation of SNr with STN when treating patients with DBS-resistant FoG, it is essential to assess and understand its mechanistic effect on cortico-subcortical networks.

The effects of DBS are thought to rely not only on the specific properties of individual neurons, but on the properties of large-scale brain networks (McIntyre & Hahn, 2010). The STN+SNr DBS efficacy in alleviating FoG is based on the notion that gait disturbances during advanced PD are associated with defective motor processing in the mesencephalic locomotor region (MLR) (Moro et al., 2010). The MLR is densely interconnected with the SNr, a major output nucleus of the basal ganglia. Thus, a faulty output of the basal ganglia system in PD might result in over-inhibition of the MLR due to GABAergic projections of the SNr and subsequent attenuation of locomotor activity (Scholten et al., 2017). In turn, functional inhibition of the STN and SNr due to DBS (Herrington, Cheng, & Eskandar, 2016) is likely to attenuate over-inhibitory SNr output and lead to restored gait and posture. To study the DBS network effects, we represent a cortico-subcortical network of regions involved in FoG episode generation as a directed signed graph (graph with the weights +1 or *−*1). The network includes the basal ganglia and brainstem regions as nodes and the excitatory or inhibitory synaptic connections as signed edges. The graph representation of the FoG network facilitates examining its topological properties and allows in-silico simulations of neural activity.

Our present goal is to introduce a mathematical framework that explains possible differences in the activity propagation along the cortico-subcortical projections due to the chosen DBS target (Figure 1). This framework aims to compare stimulation sites based on the projections affected by the PD symptom of interest and the network topological configuration. As large-scale basal ganglia and brainstem models often require extensive biological detail which may not be fully available empirically, here we focus on a more general model of excitable dynamics, the discrete three-state **SER model** (Messé, Hütt, & Hilgetag, 2018; Müller-Linow, Hilgetag, & Hütt, 2008). The letters S-E-R denote the basic node behavior of sus-ceptible (*S*) nodes becoming excited (*E*) by excited neighbors, then refractory (*R*), before turning once again susceptible, all in discrete time steps. Such basic excitable models, due to their small parameter space, allow exhaustive study of the effects of neuroanatomical network organization on network global dynamics and activity propagation (Garcia, Lesne, Hütt, & Hilgetag, 2012). The SER model is able to capture relevant spatiotemporal aspects of human brain dynamics (Haimovici, Tagliazucchi, Balenzuela, & Chialvo, 2013). Moreover, it was shown that the **functional connectivities** indicated by the SER model are comparable to those predicted by the well-established Fitzhugh-Nagumo model (Messé, Hütt, König, & Hilgetag, 2015). However, in contrast to the Fitzhugh-Nagumo model, the SER model allows to observe the entirety of all available emerging dynamical patterns of the network after initializing the graph nodes with all possible initial state conditions. After some time, the dynamic behavior of the network converges to some repeating patterns, the network attractors. A change in the graph topology, such as due to projections silenced by DBS, results in a change in the **attractor space** of the system. This attractor space, in turn, can be converted into a coactivation matrix, as a measure of functional connectivity. Thus, a change in **network topology** results, via its interpretation through the excitable model, in a change of the functional connectivity of the network.

**Figure 1.**
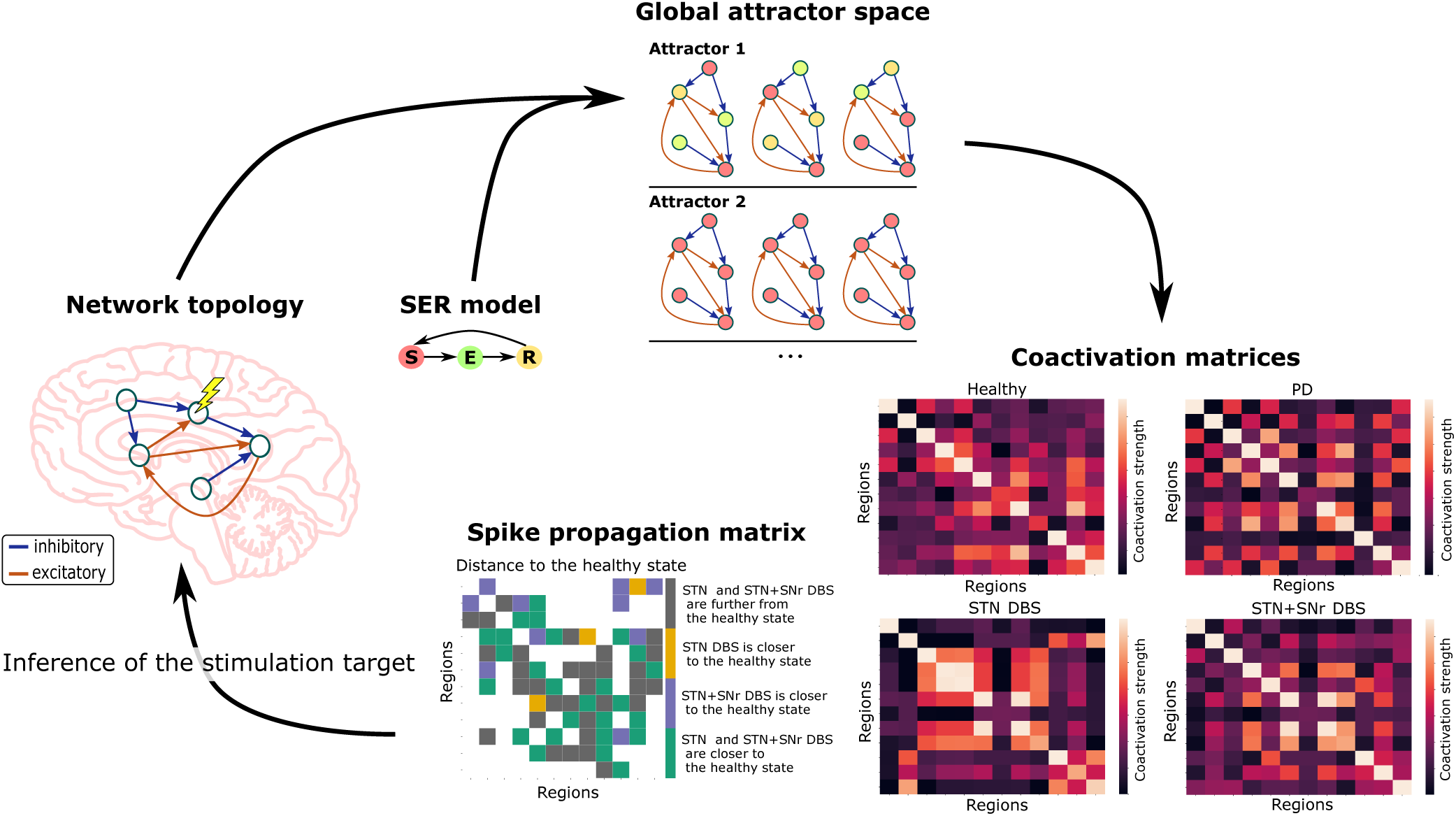
Network-based computational framework to explain the effects of targeting different DBS sites. The SER model is applied to all possible initial conditions which are available for the network nodes. This initialization leads to the emergence of different patterns of excitable dynamics, the network attractors. There are two types of attractors the system can be in: a limit cycle or a fixed point. During a limit cycle, the system repetitively goes through a set of *S* - susceptible, *E* - excited, and *R* - refractory states (attractor 1). By contrast, in the fixed point, all of the network nodes remain in the *S* state (attractor 2). The attractor space constitutes the coactivation and spike propagation matrices of the system. A change in the network topology due to a DBS target site choice affects these coactivation and spike propagation matrices. Thus, one can infer a suitable stimulation target from the changes in these matrices.

In this study, we explore the changes in dynamical landscapes emerging from the FoG network via the SER dynamics during the STN and STN+SNr DBS modes. We strive to detect the changes in activity propagation in the MLR leading to an alleviation of FoG. We hypothesize a normalization of activity propagation in the MLR due to STN+SNr DBS. A further aim of our work is to introduce a general computational network-based framework helping to elucidate large-scale effects resulting from DBS. The framework is not primarily meant to render realistic neuronal dynamics (Garcia et al., 2012), but rather to extract the essential effects of the topological changes caused by DBS on whole-brain network activity patterns.

## RESULTS

To study the changes in emerging dynamics induced by the system’s state, we employ the network of twelve regions shown in Figure 2. The network includes the basal ganglia and brainstem regions thought to be underlying FoG episode generation (Snijders et al., 2016). The FoG network can be in the healthy, PD, STN DBS, and STN+SNr DBS configurations. The PD configuration is obtained from the healthy configuration by setting the weights of all the edges originating from the SNc to 0 (five edges). The same is done for the STN DBS and STN+SNr DBS configurations based on the PD state, by setting the weights of all the edges originating from STN (eight edges), or both STN and SNr (fourteen edges), to 0. In the following sections, we first assess the dynamical patterns emerging from these network configurations and then analyze corresponding coactivation matrices. Finally, we analyze the activity propagation in different FoG network configurations. Further details can be found in the Materials and methods section.

**Figure 2.**
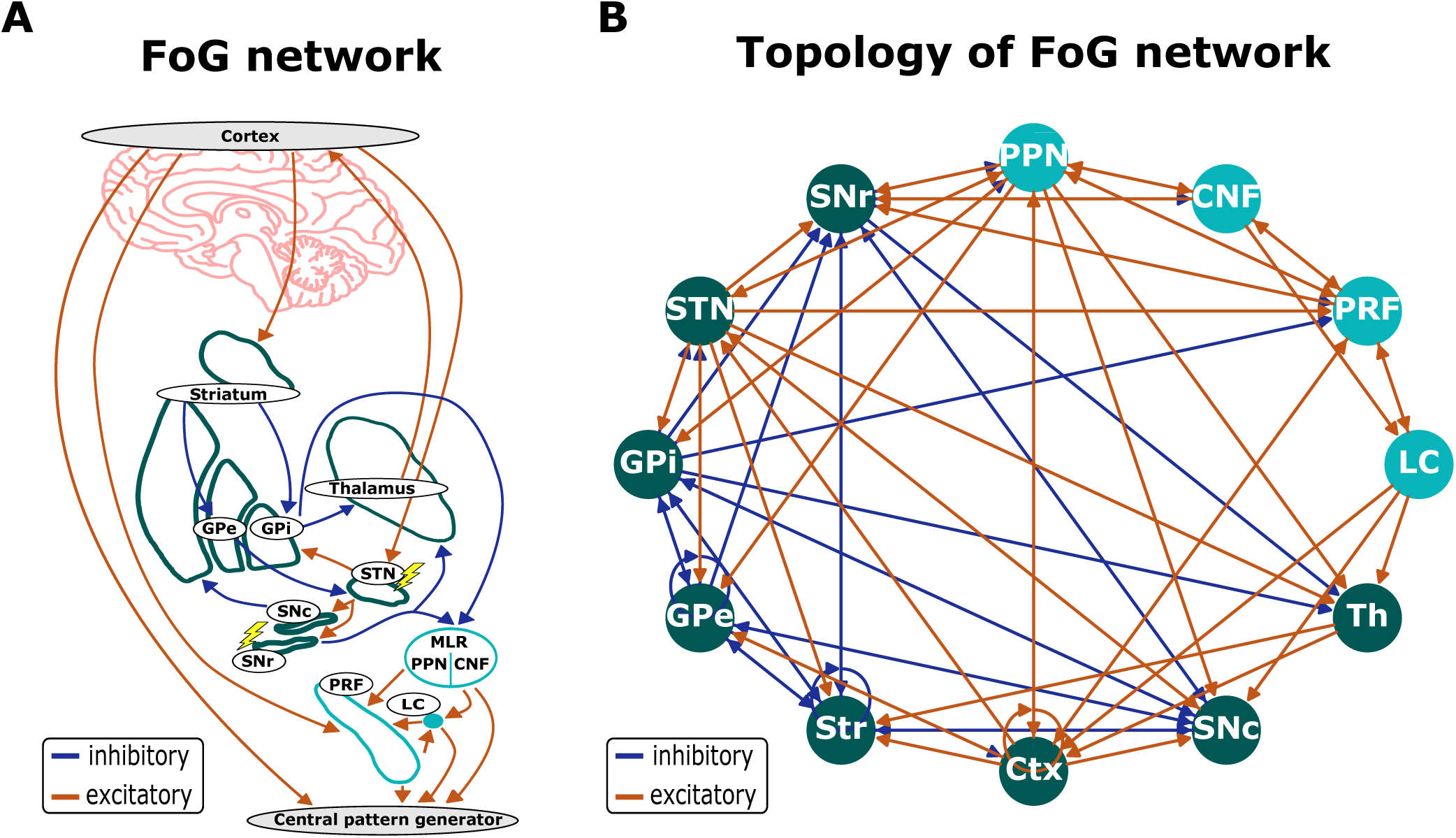
FoG network schematics. (A) Simplified schematic of the signal flow involved in gait generation and posture control. Lightning symbols indicate potential DBS targets. Central pattern generator symbolizes output neurons in a spinal cord, which are not modeled in this study. (B) Graph representation of the FoG network used in the model. Inhibitory and excitatory synaptic connections are shown as blue and red arrows, respectively. The basal ganglia regions and cortex are shown in green, whereas the brainstem regions are in turquoise. Abbreviations: Ctx - cortex, SNc - substantia nigra pars compacta, striatum, GPi - globus pallidus pars interna, GPe - globus pallidus pars externa, STN - subthalamic nucleus, Th - thalamus, SNr - substantia nigra pars reticulata, MLR - mesencephalic locomotor region, PPN - pedunculopontine nucleus, PRF - pontine reticular formation, CNF - cuneiform nucleus, LC - locus coeruleus.

### Emerging dynamical patterns

Emerging dynamical landscapes for the healthy, PD, and DBS configurations of the FoG network are derived from the excitation patterns of the deterministic SER model (see Materials and methods). These patterns regarding the FoG network configurations are summarized in Table 1. The numbers of fixed points and period-3 **limit cycles** vary largely depending on the network configuration. No limit cycles with a period other than 3 is found for the network configurations under study. It should be noted that for the configurations with a smaller number of period-3 limit cycles (healthy and STN DBS configurations), the number of unique period-3 limit cycles is also smaller.

**Table 1.**
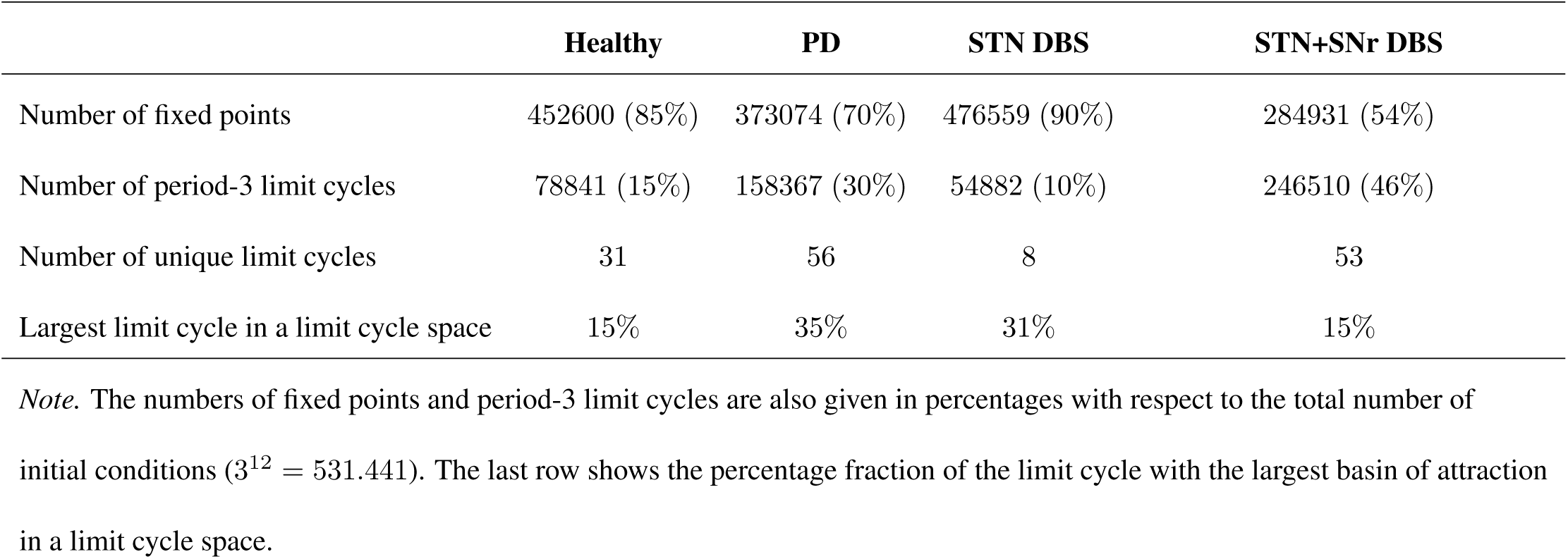
A summary of dynamical landscapes for the different FoG network configurations.

An example of an emerging dynamical landscape for the healthy network configuration is shown in Figure 3. It is notable that the largest proportion of all initial conditions converges to a fixed point (Figure 3 right inset, Table 1). In a fixed point, all the network nodes are constantly in the susceptible state. All the other initial conditions converge to 31 different period-3 limit cycles. The limit cycle with the largest basin of attraction is shown in the left inset of Figure 3. Every node in the graph repeatedly goes through a loop of *S*, *E*, and *R* states. The coactivation pattern of nodes is different for different period-3 limit cycles. Additionally, Table 1 shows that the period-3 limit cycle with the largest basin of attraction constitutes more than 30% of the limit cycle space for the PD and STN DBS network configurations.

**Figure 3.**
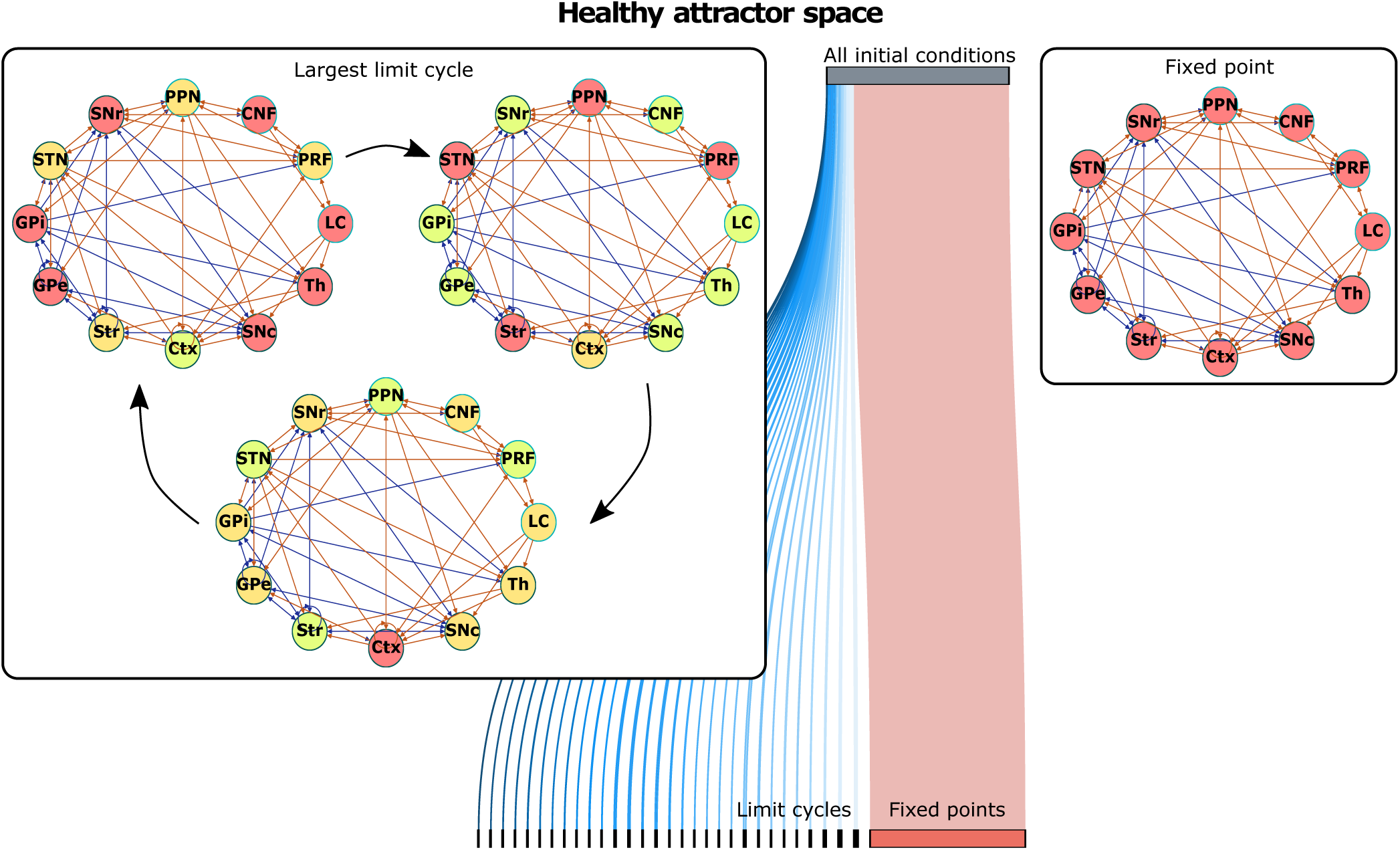
Emerging dynamical landscape for healthy network configuration. For the different combinations of the initial conditions of the SER dynamics, the healthy network topology converges to fixed points (right inset) and period-3 limit cycles (the limit cycle with the largest basin of attraction is shown in the left inset).

From an additional analysis, we note that, for the 42% of the limit cycles of the healthy network configuration, the striatum is always in a susceptible state *S*. By contrast, this is only the case for 4% of the limit cycles of the PD network configuration. For the STN DBS and the STN+SNr DBS, the proportion rises back to 17%. These proportions could be related to the firing probability of the striatum in the model. The more limit cycles exist that have a particular region always in a susceptible state, the smaller the firing probability of this particular region.

To compare the emerged limit cycles for different topological configurations of the network, we utilize Venn diagrams (Figures 4-6). We do not compare the fixed points in our subsequent analysis, as in our model a fixed point is equivalent to the absence of activity in nodes.

**Figure 4.**
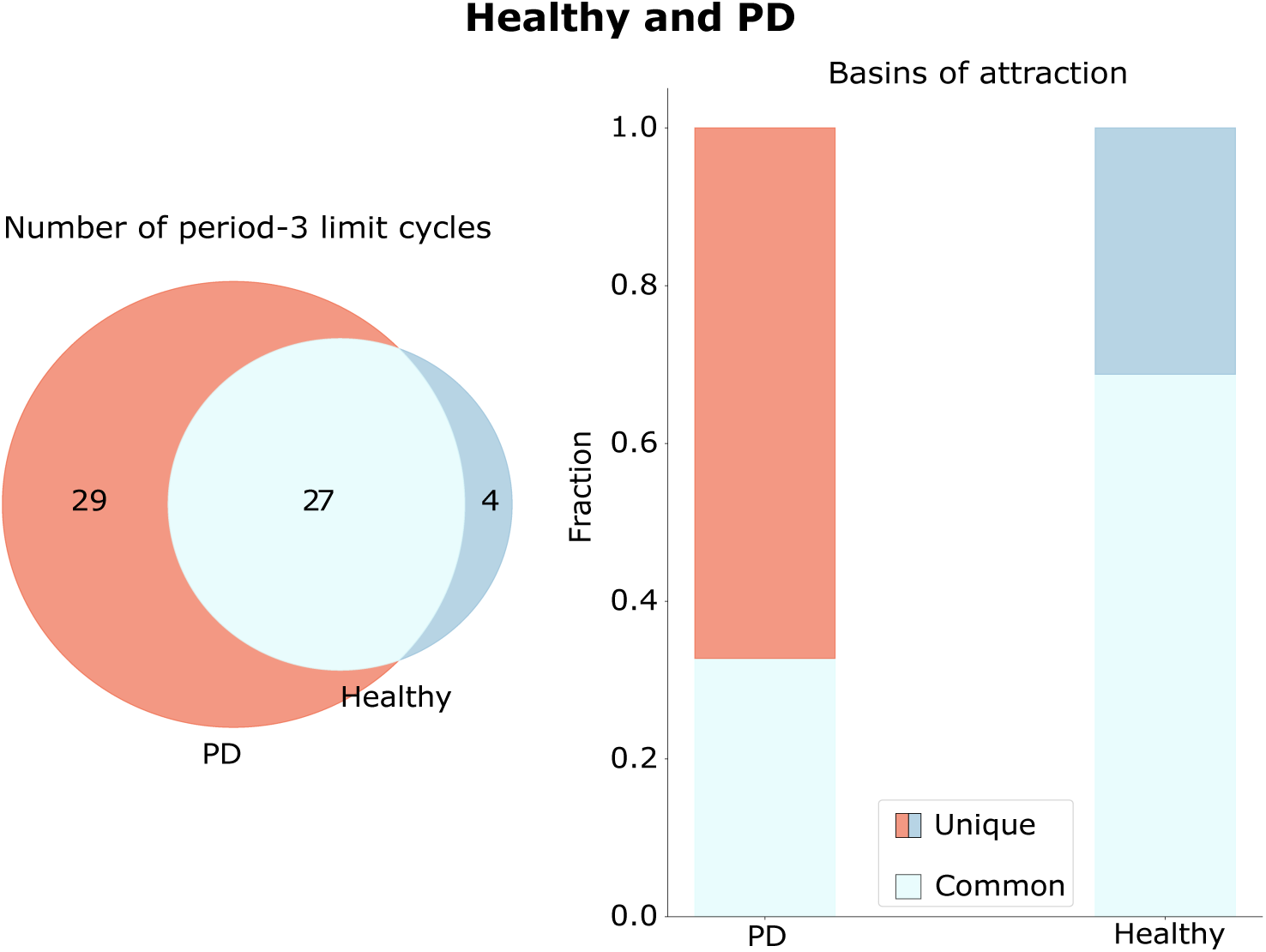
Healthy and PD limit cycle spaces. Limit cycles unique to the healthy and PD configurations are shown in blue and red, respectively. Limit cycles, which are similar in the healthy and PD configurations, are shown in light blue. The numbers inside the circles are the numbers of the limit cycles of an aforementioned type. On the right side, the basins of attractions are compared across configurations, with the fraction of the basin size to the number of limit cycles depicted along the Y-axis.

We create PD and DBS topological configurations of the FoG network by changing the weights of all edges coming from a certain node to 0 (see section Materials and methods for further details). In a deterministic SER model, this approach is equivalent to an external control exerted over the healthy, or, in the DBS case, PD FoG network configuration (Borriello & Daniels, 2020). When an external control is exerted on a network, the basin of original attractors of the network can change, or new attractors can emerge (Newby, Tejeda Zañudo, & Albert, 2022). Thus, a successful intervention, in our case DBS, can be viewed as the steering of the PD system towards an attractor space of the original healthy system.

The attractor spaces of the healthy and PD configurations are compared in Figure 4. It can be seen that, during the exerted external control of moving from the healthy to the PD configuration, new attractors emerge (red attractors in Figure 4). In addition, the basins of attraction of the original attractors change (right panel in Figure 4). Moreover, it is noticeable that healthy and PD configurations share several common limit cycles. An additional analysis shows that the limit cycle with the largest basins of attraction in the PD configuration does not appear in the healthy configuration. From the right panel in Figure 4, one can see that the fraction of the basins of attraction of the unique healthy limit cycles is much smaller than the fraction of the ones that appear in both the healthy and the PD configurations. Thus, DBS is less likely to steer the system back towards its unique attractors of the healthy configuration than to the attractors shared in common between the configurations, or to some new attractors.

In Figure 5, the limit cycle spaces of the healthy, the PD, and the STN DBS configurations are compared. One can see that only two new attractors are not present in the attractor space of the PD or healthy configurations (in yellow). At the same time, the dynamical landscape of the STN DBS network includes one of the attractors of the healthy configuration, which are not the same as for the PD state (in dark blue). This attractor has the striatum node always in the *S* state, which again points out the low spiking activity of the striatal neurons in the healthy conditions (Singh et al., 2016), as all the other nodes in the attractor are going through the sequence of *S*, *E*, and *R* states. From an additional analysis, we find that in the STN DBS configuration, the system moves away from the limit cycle with the largest basin of attraction in the PD configuration. On the right panel in Figure 5, the basins of attraction of different attractor groups are compared. The fraction of unique attractors of the STN DBS configuration (in yellow) and the attractor present in both STN DBS and the healthy configuration (dark blue) is small compared to the other types.

**Figure 5.**
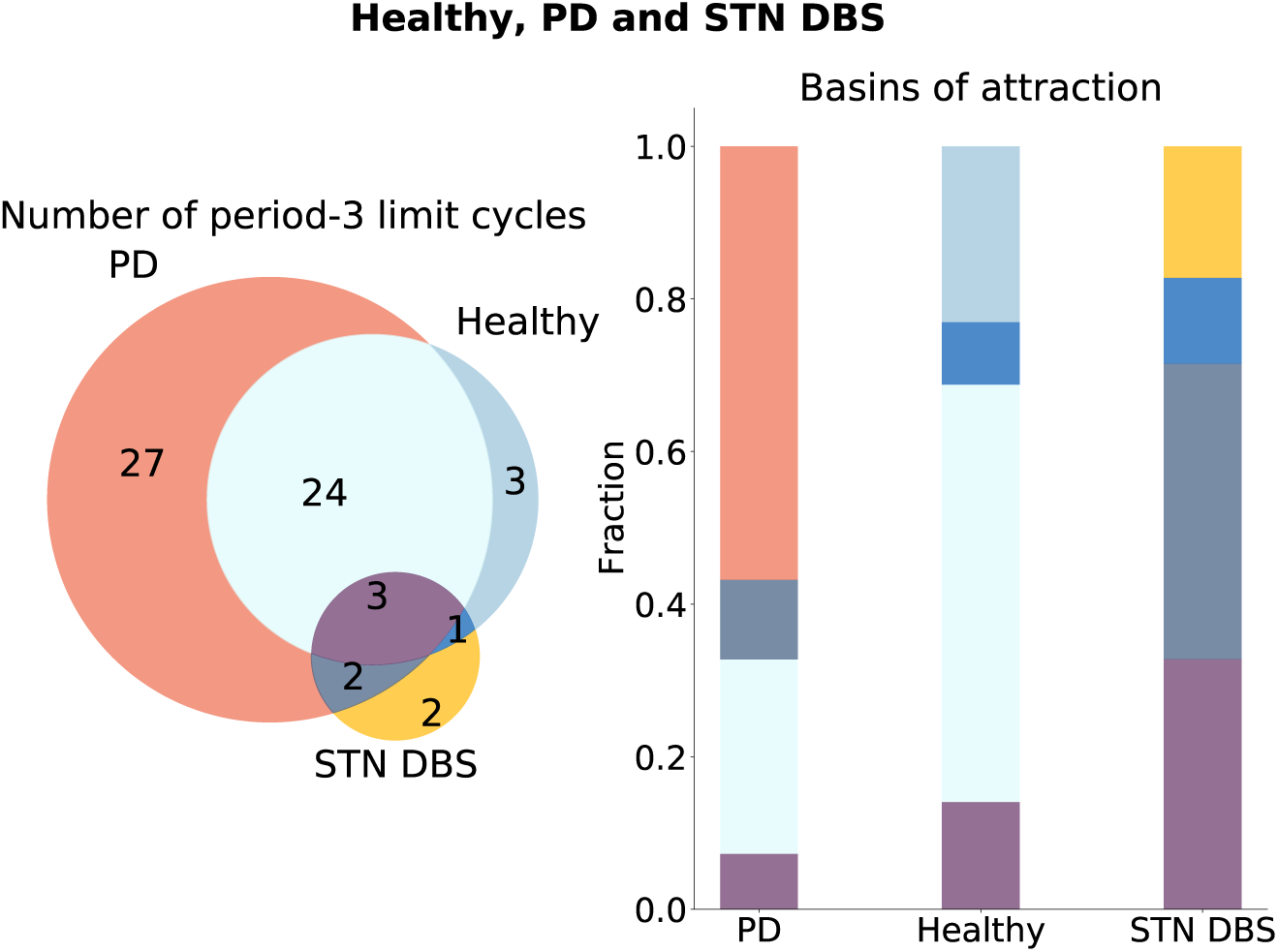
Healthy, PD, and STN DBS limit cycle spaces. Limit cycles unique to the healthy, PD, and STN DBS configurations are shown in blue, red, and yellow, respectively. Limit cycles, which are similar in the healthy and PD configurations, are shown in light blue. Limit cycles, which are similar in the STN DBS and PD configurations, are shown in gray. Limit cycles, similar in the STN DBS and healthy configurations, are shown in dark blue. Limit cycles, which are similar across all configurations, are shown in purple. The numbers inside the circles are the numbers of the limit cycles of an aforementioned type. On the right side, the basins of attractions are compared across configurations, with the fraction of the basin size to the number of limit cycles depicted along the Y-axis. The colors correspond to the colors on the left panel.

In Figure 6, the limit cycle spaces of the healthy, the PD, and the STN+SNr DBS configurations are compared. In contrast to the STN DBS, one can see that many new attractors appear that are not present in the attractor space of the PD or healthy configurations (in yellow). At the same time, the dynamical landscape of the STN+SNr DBS network includes one unique attractor of the healthy configuration (in dark blue). This is the same attractor as in Figure 5. However, the whole system rather moves into the new dynamical space, as there are 36 new limit cycles, which constitute 66% of all the limit cycles in the STN+SNr DBS attractor space. The same pattern could be seen when comparing basins of attraction (right side of Figure 6). The basin of attraction of unique attractors of the STN+SNr DBS configuration (in yellow) is the largest compared to the other groups. From an additional analysis, we find that in the STN+SNr DBS configuration, the network also moves away from the limit cycle with the largest basin of attraction in the PD configuration.

**Figure 6.**
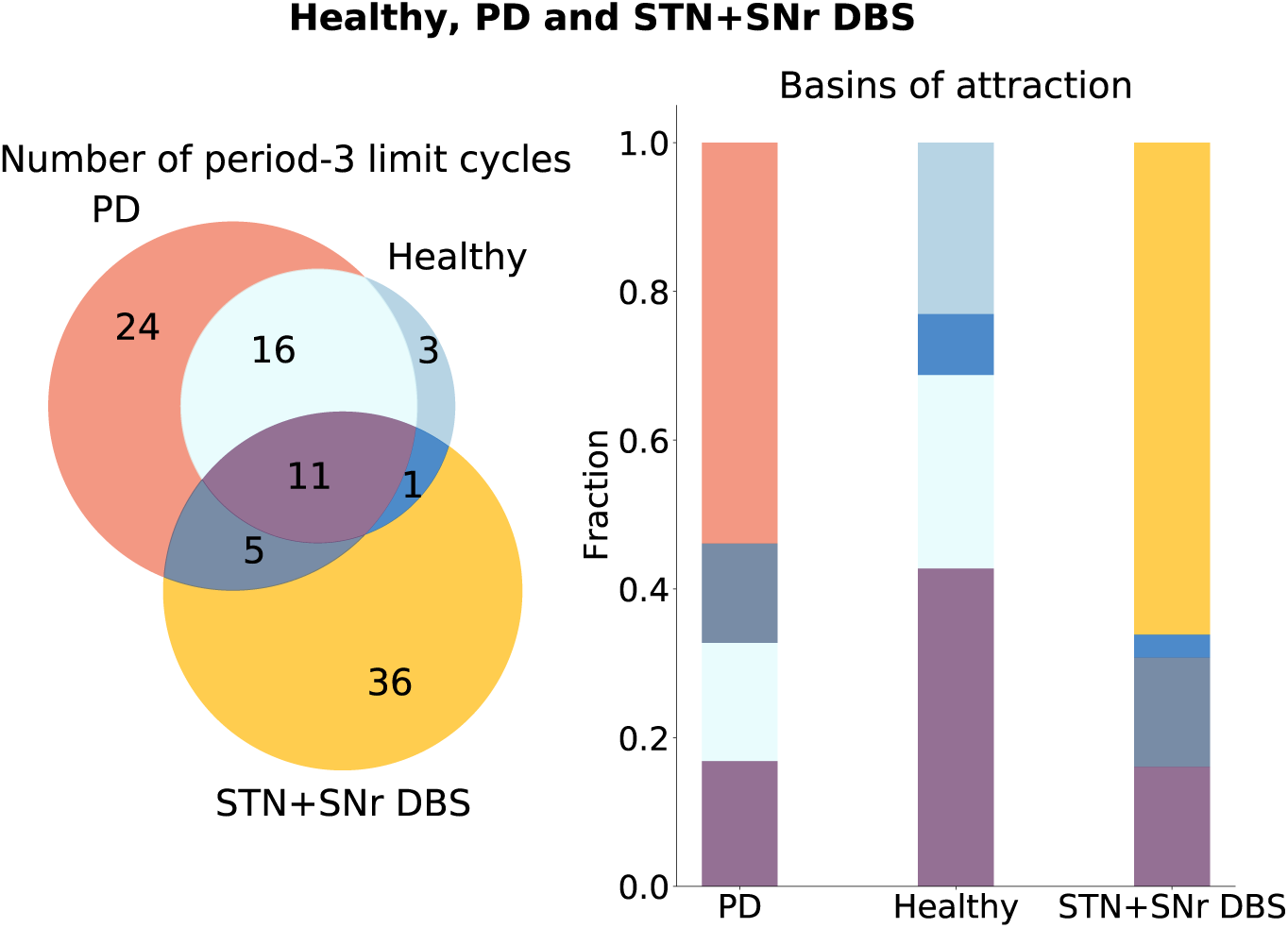
Healthy, PD, and STN+SNr DBS limit cycle spaces. Limit cycles unique to the healthy, PD, and STN+SNr DBS configurations are shown in blue, red, and yellow, respectively. Limit cycles, which are similar in the healthy and PD configurations, are shown in light blue. Limit cycles, which are similar in the STN+SNr DBS and PD configurations, are shown in gray. Limit cycles, similar in the STN+SNr DBS and healthy configurations, are shown in dark blue. Limit cycles, which are similar across all configurations, are shown in purple. The numbers inside the circles are the numbers of the limit cycles of an aforementioned type. On the right side, the basins of attractions are compared across configurations, with the fraction of the basin size to the number of limit cycles depicted along the Y-axis. The colors correspond to the colors on the left panel.

### Coactivation matrices

The coactivation matrices for the healthy, the PD, and the DBS configurations are shown in Figure 7. The details behind the implementation of the coactivation matrices can be found in the Materials and methods section. The coactivation matrices depict the level of synchronous activity (*i.e.,* a measure of functional connectivity) of the different regions across all limit cycles. The larger the coactivation strength, the more synchronously the two regions are firing on average.

**Figure 7.**
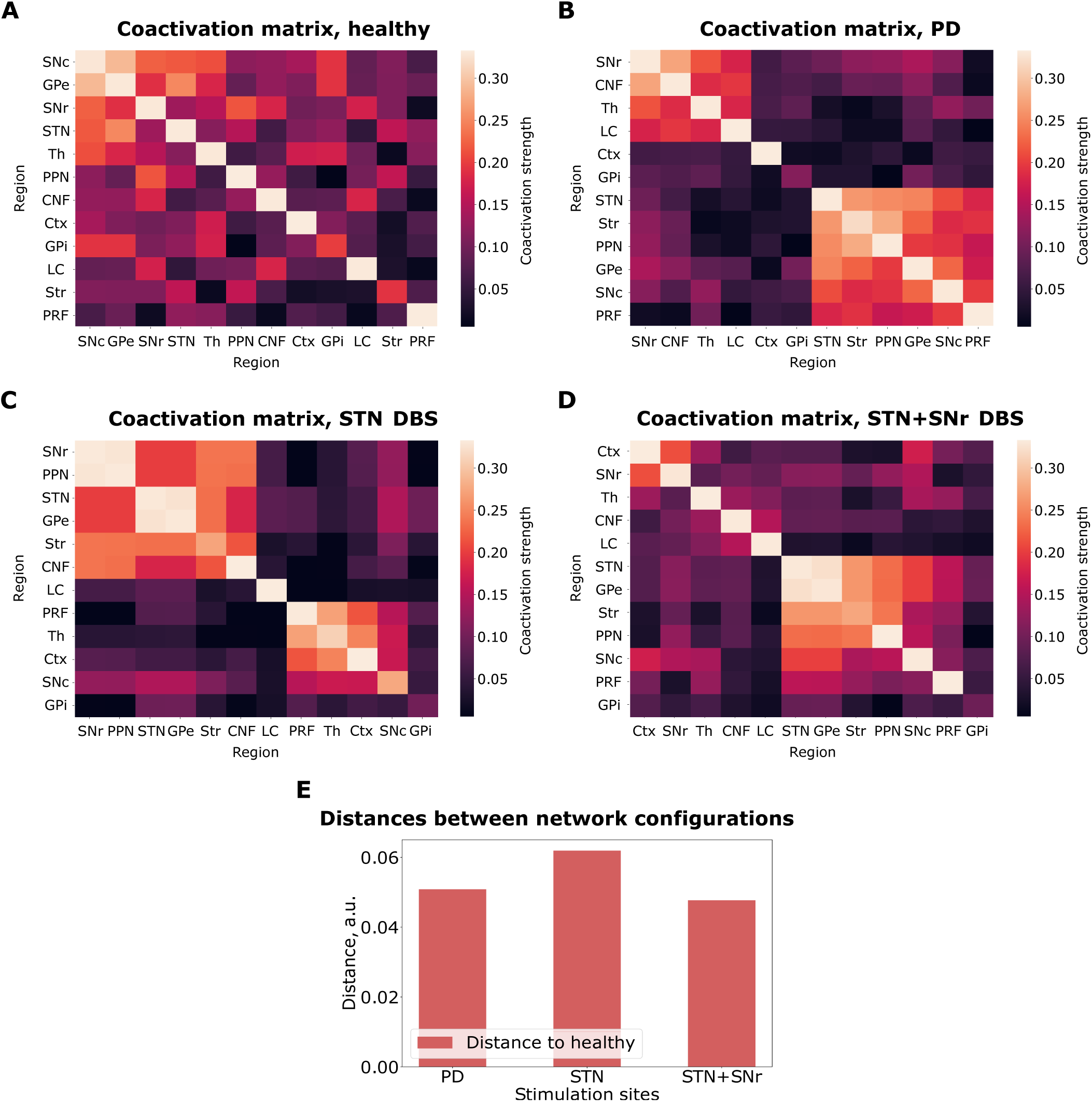
Coactivation matrices. (A) The healthy configuration. (B) The PD configuration. (C) The STN DBS configuration. (D) The STN+SNr DBS configuration. (E) 1-norm distances between the coactivation matrices of the PD, STN DBS and STN+SNr DBS to the healthy state.

The coactivation matrices in the healthy and the PD states differ (Figure 7 A, B). The pattern of activity in the healthy matrix appears to be smooth, whereas the pattern in the PD coactivation matrix appears to contain modules. The modular structure is also visible in the STN DBS coactivation matrix (Figure 7 C). However, the regions participating in the co-active groups are different from the ones that participate in the co-active groups in the PD configuration (Figure 7 B). In the STN+SNr DBS case, the modular organization appears to dissipate. Figure 7 E shows how close the DBS coactivation matrices are to the healthy one. We evaluated the similarity by using the 1-norm distance. Indeed, the STN+SNr DBS coactivation matrix is more similar to the healthy state coactivation matrix relative to the PD configuration. On the other hand, the STN DBS coactivation matrix does not exhibit this behavior.

### Activity propagation

There is a hypothesis that the normalization of excitatory signals propagation in brainstem regions might be the reason for successful STN+SNr DBS during FoG (Pötter et al., 2008). Thus, we assess not only the general coactivation activity of the regions, but also the activity-based projections between them. To this end, we use the activity spike propagation matrix (Figure 8). The details behind its implementation can be found in the Materials and methods section.

**Figure 8.**
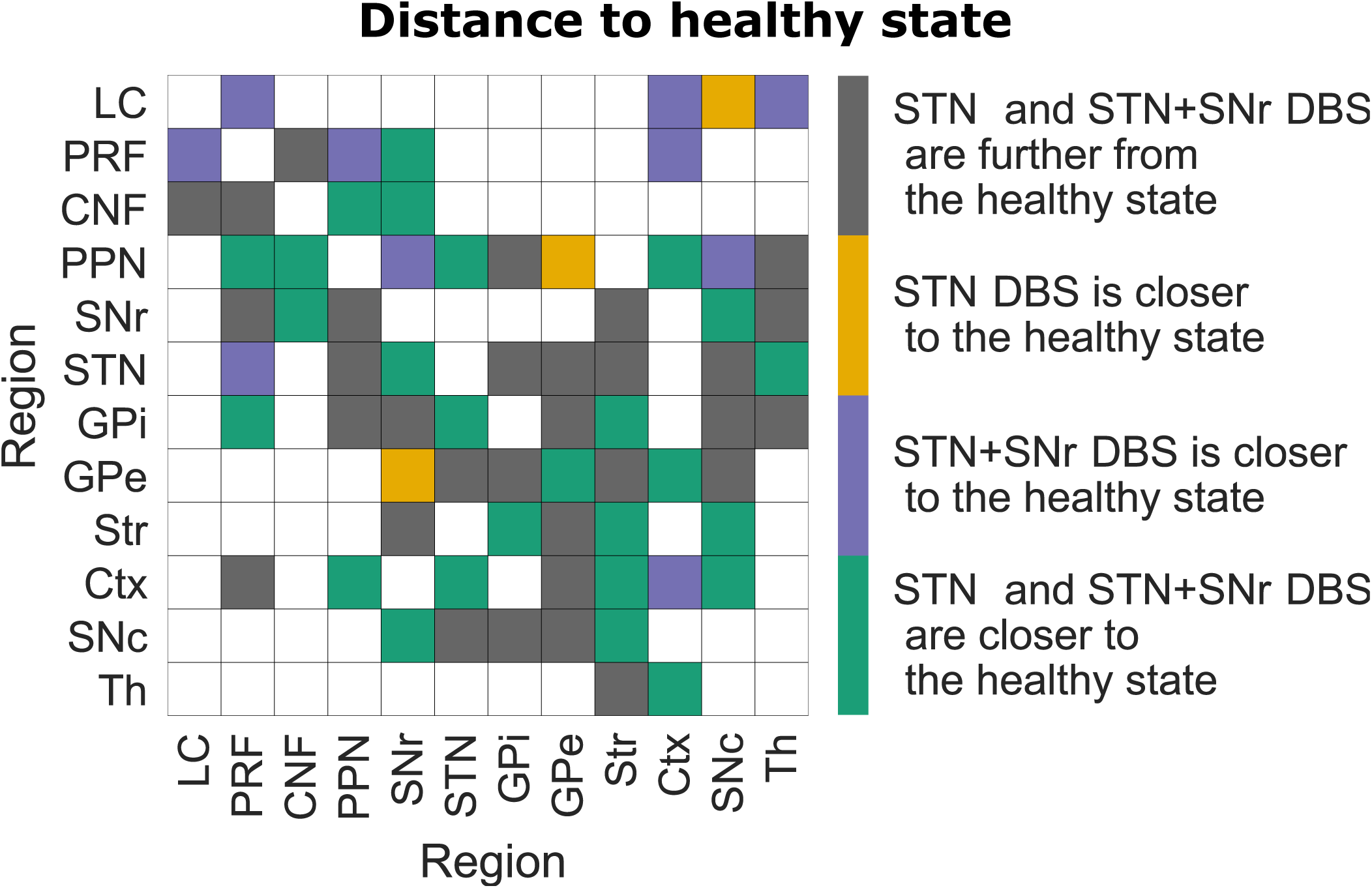
Spike propagation flow normalization. Projection source regions are shown along the Y-axis, while projection targets are shown along the X-axis. The projections are color-coded based on how far they are from the healthy state for different DBS configurations compared to the distance between the healthy and PD configurations. Being far is equivalent to having a larger 1-norm distance between the elements in the coactivation matrix for excitatory and inhibitory connections between the healthy and DBS configurations than between the healthy and PD configurations (implementation can be found in the Materials and methods section).

Figure 8 shows a general normalization of the spike flow during the STN+SNr DBS configuration (31 gray squares, 3 yellow squares, 10 purple squares, 26 green squares). However, it is also noticeable that for the normalization of excitatory propagation along the brainstem projections, the STN+SNr DBS is exclusively effective (purple squares for the LC-PRF, PRF-LC, PRF-PPN, LC-Ctx projections). Figure 8 also reveals that STN DBS is more beneficial for specific projections (yellow squares). This observation indicates that, for these projections, the signal propagation is closer to the healthy configuration during the STN DBS configuration than the distance in signal propagation in these projections between the healthy and the PD configuration. In the case of yellow projections, it means the opposite for the STN+SNr DBS configuration. These yellow projections could be the projections of concern for the STN+SNr DBS configuration.

## DISCUSSION

In this study, we present a new network-based computational framework to uncover dynamical landscapes and activity propagation in the brain networks relevant for understanding symptoms of Parkinson’s disease. The framework aims to compare DBS targets depending on the projections of importance. In the context of FoG in PD, we found that STN+SNr DBS outperforms the standard STN DBS in terms of activity propagation normalization in the brainstem regions (Table 2). Activity propagation flow is closer to the healthy state propagation flow in LC-PRF, PRF-LC, PRF-PPN, and LC-Ctx projections for the STN+SNr DBS. This result aligns with previous clinical studies which found that STN+SNr DBS outperforms STN

**Table 2.** A summary of study hypotheses.

| Hypothesis | Corroborated | Contradicted |
| --- | --- | --- |
| Under the DBS configurations the network attractor space is similar to the healthy attractor space |  | ✓ |
| Under the DBS configuration the network attractor space rather moves to the new attractor space | ✓ |  |
| STN DBS normalizes spike propagation flow in the FoG network |  | ✓ |
| STN+SNr DBS normalizes spike propagation flow in the FoG network | ✓ |  |
| STN+SNr DBS outperforms STN DBS in normalizing spike propagation flow in the FoG network | ✓ |  |
| STN+SNr DBS normalizes spike propagation flow in the brainstem projections | ✓ |  |

DBS in the context of FoG symptoms (M. A. Horn et al., 2022; Wagner et al., 2022; Weiss et al., 2013). According to the hypothesis in (Scholten et al., 2017), SNr stimulation leads to temporal regularization of gait and normalization of the output to the MLR and brainstem projections signaling, which is corroborated by the present results. On the other hand, our model suggest that STN DBS does not normalize spike propagation flow in the FoG network (Table 2). This observation is at odds with STN DBS usually improving patient symptoms (Deuschl et al., 2013). We believe that the discrepancy is due to the choice of network topology. In the current study we were particularly focusing on the FoG network, whereas to study general symptom normalization in the course of the PD one probably should refer to more specific tremor (Duval, Daneault, Hutchison, & Sadikot, 2016), dystonia or bradykinesia networks.

To obtain the present results, we first checked dynamical landscapes of the FoG network in healthy, PD, STN DBS, and STN+SNr DBS configurations using the discrete excitable SER model. The model approach might appear simplistic, although it exhibits the most important properties of the three-state excitatory cycle of neural elements. At the same time, the amount of biological realism required for an optimal spiking model to capture the PD dynamics is an open question. Our network-based SER framework is not intended to render realistic neuronal firing patterns. Nevertheless, it captures essential aspects of the PD and DBS dynamics. Analysis of attractors with the largest basins of attraction (Table 1) reveals that, for the PD FoG network configuration, the largest basin of attraction constitutes 35% of the limit cycle space. This is a large increase compared to the healthy configuration (15% of limit cycle space). The increase might be interpreted as a sign of abnormal synchrony within the basal ganglia nuclei during PD (Wichmann & DeLong, 2006). During the STN DBS, the amount of limit cycle space taken up by the attractor with the largest basin of attraction does not fall significantly (31%). This finding might indicate the topological change leading to STN DBS regularization of the firing patterns in basal ganglia (McConnell, So, Hilliard, Lopomo, & Grill, 2012). Indeed, the activities of basal ganglia nuclei are highly correlated with one another for the attractor with the largest basin of attraction in the STN DBS configuration (Figure S1). Additionally, there is a change in activity of the striatal neurons. The number of striatal neurons in a susceptible state *S* decreases in the PD topology, when compared to the healthy topology. Striatal neurons are known to be nearly quiescent in the healthy condition, while their firing rates rise to about 17 Hz during the PD (Singh et al., 2016). However, the human recordings by Moll et al., 2015 (Moll, Hamel, & Engel, 2015) suggest the PD value to be 2 *−* 5 Hz. During DBS, the firing rate of striatal neurons tends to go back to the healthy range independently of the chosen DBS target.

The network-based SER framework allows exhaustive assessment and comparison of the emerging dynamical landscapes (Figures 4, 5, 6). This approach helps to address the working mechanism of DBS at the network level. We can see that the system moves away from the most prominent attractors in the PD configuration for both DBS conditions (Figures 5, 6). Although moving towards the attractors of the healthy configuration is less likely (Figure 4, right panel), both DBS attractor spaces incorporate one unique attractor of the healthy configuration (Figures 5, 6). Otherwise, the system for the DBS exhibits a trend to move away to a new attractor space (yellow parts of Figures 5, 6, Table 2). This observation is in accordance with the conclusions by Wichmann and DeLong (Wichmann & DeLong, 2006). They state that the system for the DBS reaches a new equilibrium, rather than induces restoration of normal basal ganglia function. We continued to study the new equilibrium DBS states by exploring the coactivation matrices and spike propagation flow in the FoG network. The coactivation matrices revealed that combined STN+SNr DBS is closer to the healthy state in terms of functional connectivity (Figure 7). Furthermore, the pattern of activations in the PD coactivation matrix appears to contain modules, which affirmed observed patterns of synchronous activity in basal ganglia nuclei (Hammond, Bergman, & Brown, 2007; Magnin, Morel, & Jeanmonod, 2000). The spike propagation flow normalization in the MLR (Figure 8) exclusive to the STN+SNr DBS reaffirms the hypothesis on its working mechanism by (Scholten et al., 2017). Additionally, it captures the normalization of the cortico-thalamic connectivity towards the healthy controls for DBS (A. Horn et al., 2019).

The healthy FoG network topological configuration governs the resulting dynamical output, as network topology is the most influential parameter in the SER model. In the present study, we used the topology described in Figure 2. The initial choice of this signed graph heavily affects the observed dynamical patterns. We motivated this network choice by comparing the healthy FoG network configuration with a mesoscale connectome from (Oh et al., 2014) (Figure S2). Specifically, we compared the weights of the edges from the Allen brain atlas data in (Oh et al., 2014) with the corresponding weights of the edges in the healthy FoG network in Figure S2. As the weights in (Oh et al., 2014) correspond to the tract-tracing data, they are strictly non-negative. That is why we assign the sign of the efferent projections from the cholinergic PRF, PPN, CNF, and LC based on (Pahapill, 2000; Snijders et al., 2016). The basal ganglia region’s sign and projections are based on (Fleming, Dunn, & Lowery, 2020; Lourens, Meijer, Heida, Marani, & Gils, 2011; So, Kent, & Grill, 2012). From the Figure S2, it can be seen that the regions without edge connections in the FoG network correspond to the group of edges with the weights clustered around 0 (orange violin plot). Otherwise, strong inhibitory or excitatory connections cluster around *−*1 and 1 in the FoG network, respectively, as expected. There are no edges with large weights in (Oh et al., 2014), which would correspond to a 0 weight in the FoG network. Thus, the healthy FoG network configuration in Figure 2 depicts the most significant connections in the Allen brain atlas connectome (Oh et al., 2014), which supports the reliability of the chosen topology. However, a strong reliance on a choice of the topology is still one of the limitations of the current approach. That is why we propose to use the current modeling framework after carefully assessing the neuroanatomy behind a symptom under study.

Another related limitation of the current computational framework is its computational load. The amount of initial conditions needed to be assessed to determine the attractor space of the network scales exponentially with the network size. That is why we have opted to a FoG network of a smaller size and have not included further brain networks, such as the limbic system, in the present study.

Overall, we propose a practical computational framework that simultaneously captures the essential aspects of PD and DBS, explores the new STN+SNr DBS approach and validates its efficiency in the context of FoG.

## CONCLUSION

Selecting the best therapeutic strategy for a specific symptom in PD can be burdensome. In this study, we propose a straightforward network-based computational framework to determine a suitable DBS target for treating FoG. This approach is based on a basic discrete SER model and requires only general knowledge of the cortico-subcortical network topology. We also showed that STN+SNr DBS may be more beneficial for patients with FoG due to the normalization of the spike propagation flow in the MLR, which confirms our initial hypothesis. In the future, we aim to explore additional DBS targets and network regions, specifically in the limbic system, to investigate the influence of STN+SNr DBS on mood and anxiety.

## MATERIALS AND METHODS

### The FoG network

The freezing of gait network (Figure 2) was used to compare possible network effects induced by STN DBS and STN+SNr DBS. These effects were then used to identify the possible working mechanisms behind DBS and to infer a suitable stimulation mode for FoG. The network consists of the basal ganglia, the motor cortex, the thalamus, the pedunculopontine nucleus, the pontine reticular formation, the cuneiform nucleus, and the locus coeruleus. The choice of the network regions and connections was based on an analysis of the anatomical literature (Pahapill, 2000; Snijders et al., 2016; Takakusaki, 2017). A schematic for locomotion control (Snijders et al., 2016) was a primary resource used for the construction of the network. The brainstem circuitry (PRF, CNF) was also partially taken from the Mouse Brain Connectivity Atlas provided by Allen Institute (Allen Institute for Brain Science (2004), 2011; Oh et al., 2014) in order to expand the network. We assigned the sign of the efferent connections from the PRF, CNF and LC based on (Snijders et al., 2016) assuming homogeneity of the neuromodulator type across the connections. The basal ganglia and PPN connections were largely taken from (Guatteo, Cucchiaroni, & Mercuri, 2009; Lourens et al., 2011; Pahapill, 2000; So et al., 2012).

The FoG network can be represented as a directed signed graph. The nodes of this graph are the brain regions which are involved in the FoG network. The weights of the edges are set to +1 or *−*1 in the case of the excitatory and inhibitory synaptic projections, respectively. To compare the possible effects induced by STN or STN+SNr DBS on the FoG network, we explored their dynamics in four possible *configurations*: healthy, PD, STN DBS, and STN+SNr DBS. To implement the PD configuration, we set the weights of all edges originating from the SNc node to 0. This approach was chosen because Parkinson’s disease is characterized by the degeneration of dopaminergic neurons in SNc, which results in a dopamine depletion in the efferent targets of the SNc (Delaville, de Deurwaerdère, & Benazzouz, 2011). In our model, this degeneration was taken to be equivalent to the disconnection of the SNc from the FoG network. The DBS configurations were created from the PD by setting the weights of all the edges originating from the stimulation target node to 0. This approach is in accordance with the virtual lesion hypothesis being the working mechanism behind the DBS (Herrington et al., 2016). The hypothesis is based on the similarity between the effects observed after surgical lesions of the brain regions and their high-frequency DBS. According to this idea, DBS is thought to function via inhibition of the neurons in the vicinity of the stimulating electrode. In our model, this phenomenon was taken to be equivalent to the disconnection of the DBS target region. Thus, in the case of STN DBS, the weights of all the edges originating from the STN node were set to 0. During the combined STN+SNr DBS, the weights of the edges originating from both the STN node and the SNr node were set to 0.

### The SER model

We studied all possible dynamical patterns which emerge in the healthy, PD, and DBS FoG networks. Thus, for computational effort, a minimal excitable dynamical system, a discrete excitable SER model (Messé et al., 2018; Müller-Linow et al., 2008) was used. In the SER model, the *S* stands for susceptible, *E* excited, and *R* refractory states. In the deterministic version of the model, the node goes through the aforementioned states. If the node is in a susceptible state *S* and if the sum of the weights of the edges originating from excited neighbouring nodes and converging on the target note is larger than 0, then in the following discrete time step, this node will become excited (*E*). From the *E* state, the node will always go to a refractory state *R*. In turn, from the *R* state, the node will always go to the state *S* in the following discrete time step.

We applied the SER model over all possible initial conditions of the FoG network for the healthy, PD, and DBS network configurations to explore and compare emerging dynamical patterns. At the beginning, every node of the FoG network could be in one of the states: *S*, *E*, or *R*. After considering all possible combinations of these initial conditions, every possible dynamical pattern which can emerge in the FoG network for a certain topological configuration was explored. After that, the dynamical landscapes were compared between the healthy, PD, STN DBS, and STN+SNr DBS FoG network configurations.

As the SER model is a three-state model, there are 3*^n^* possible patterns of activity in a network with *n* nodes. These patterns of activity represent the *states* of the network. Thus, the network dynamics can be represented as a time series of the states. As the amount of the states is finite for a system with a finite number of nodes, the dynamics of the network eventually converges to one of two types of behavior, the *attractors*. The first type of attractor is a *fixed point*. The system reaches a fixed point after some transient time when all nodes in the network remain in the state *S*. The other attractor type is a *limit cycle*. In this case, all the nodes in the network will go through a series of the *S*, *E*, and *R* states with a certain period. In principle, our method can detect limit cycles with any given period. However, in the case of our networks, the detected period for all of the limit cycles was 3. Attractors could be compared via their *basin of attraction*, which is a set of initial states that converge onto a particular attractor (Borriello & Daniels, 2020). Thus, to compare the dynamical landscapes in the various FoG network states, we compared the emerging attractors and their basins of attraction for different FoG network configurations and the SER dynamics after the time *T* = 100 with the transient time *t* = 40.

Additionally, we studied coactivation and activity propagation patterns of the network in different configurations. Coactivation matrices were obtained from the time series of the network states by summing up simultaneously occurring spiking events (*E* states) across all limit cycles and dividing this sum by the simulation time. In this case, the matrix diagonal will go to the maximum of 0.33 as in a cycle with a period 3, the *E* state can happen only on every third time-step. The element at the matrix diagonal could be also less than 0.33, as in some of the limit cycles some of the nodes could always be in the state *S*.

For visualization purposes, the elements in the coactivation matrices were reordered to highlight their modular structure. For this, we used the Louvain community detection method implemented in the Brain Connectivity toolbox (Blondel, Guillaume, Lambiotte, & Lefebvre, 2008; Rubinov & Sporns, 2010). Thus, the matrix is reordered to maximize the weights and the number of edges within groups and minimize the number of edges across groups (Rubinov & Sporns, 2010). The reordering highlights groups of regions that are co-active with one another.

To calculate the activity propagation flow in the FoG network under different topological configurations, we calculate 1-norm distances between the elements of coactivation matrices corresponding to the FoG network projections. We assessed the activity propagation flow differently depending on whether the projection in the FoG network was excitatory or inhibitory.

To study excitatory projections, we used the shifted version of the coactivation matrices (Figure 9 A). It is obtained by shifting the activity of the projection target region one time step further from the activity source region. This way, if the excitatory state *E* co-occurs in the projection target and the source region, it means that the excitation of the target follows the excitation of the source. In our model, this was the case for the excitatory projections. We sum up the co-occuring shifted *E* events across all limit cycles and divide this sum by the simulation time. This way, we get a shifted coactivation matrix the elements of which we use for the excitatory projections.

**Figure 9.**
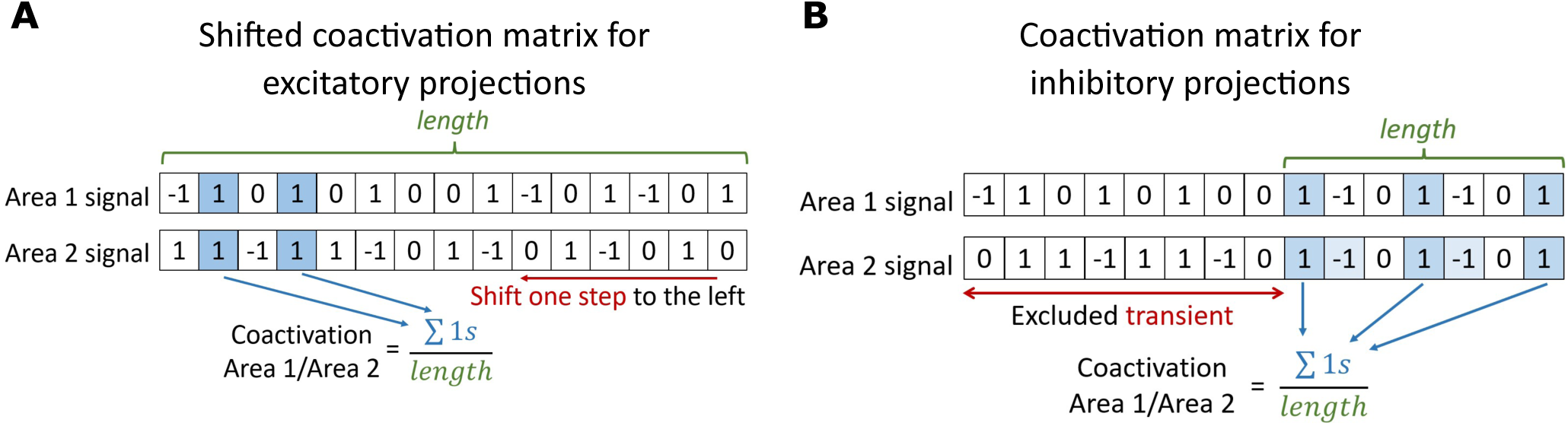
Mechanism to assess spike propagation flow. (A) Shifted coactivation matrix for excitatory projections. After the shift of one signal to the left, the co-occuring *E* states are calculated. (B) Coactivation matrix for inhibitory projections. After exclusion of transient time *t*, the co-occuring *E* states are calculated.

To assess activity propagation flow along the inhibitory projections of the FoG network, we used the usual coactivation matrices (Figure 9 B). If the excitatory state *E* occurred simultaneously in the projection target and the projection source region, it meant that the excitation of the source was followed by the target in the refractory state *R* on the next time step. This would be true only after the transient time when the system has reached its stable dynamical pattern. In our model, the excitation of the source followed by the target region entering the refractory state was equivalent to inhibition. Thus, we utilized the elements of coactivation matrices to assess the spike propagation along the inhibitory projections.

After obtaining the coactivation matrices for excitatory and inhibitory projections for every model configuration, we calculated 1-norm distances between them for each FoG network region pair and colorcoded them based on whether the distance was closer to the healthy or the PD state under different DBS configurations than the distance between the healthy and the PD configurations. This way, we obtain a matrix, an element of which shows if the 1-norm distance between the DBS and healthy configurations is smaller than the 1-norm distance between the healthy and the PD configuration. The obtained matrix was masked to depict only inhibitory and excitatory projections that exist in the FoG network (Figure 8).

## ACKNOWLEDGMENTS

MP, AM, CCH, MPN, AG, CG were funded by the Deutsche Forschungsgemeinschaft (DFG, German Research Foundation) - SFB 936 - Project-ID 178316478-A1/C1/C8/Z3.

## TECHNICAL TERMS

**Susceptible-excited-refractory model** a minimal model of excitable dynamics describing discrete signal propagation through the network.

**Deep brain stimulation** a treatment strategy for patients with Parkinson’s disease that involves electrical stimulation of brain regions.

**Functional connectivity** a statistical dependence between signals in different brain regions.

**Network topology** a set of structural connections between network nodes.

**Attractor space** a set of all possible patterns of activity in the network.

**Limit cycle** a repeating network activity pattern.

