## Supplement for "The effect of deep brain stimulation on cortico-subcortical networks in Parkinson’s disease patients with freezing of gait: Exhaustive exploration of a basic model"

*Journal:* **NETWORK NEUROSCIENCE**

RESEARCH

This document contains supplementary figures to the article entitled "The effect of deep brain stimulation on cortico-subcortical networks in Parkinson's disease patients with freezing of gait: Exhaustive exploration of a basic model".

Link to the repository: https://github.com/mariiapopova/FOG-SER

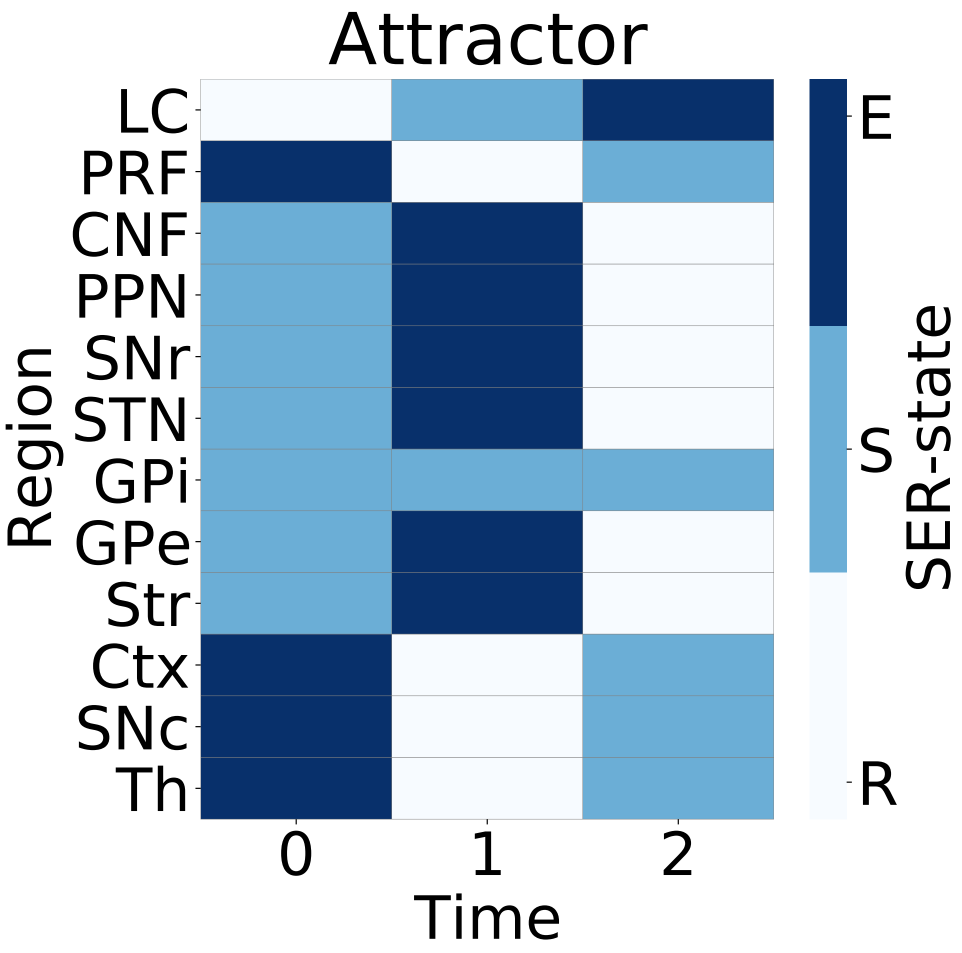


**Figure S1. Limit cycle with the largest basin of attraction for the STN DBS configuration.**

Y-axis shows different regions of the FoG network, while X-axis shows time steps. States of the SER model are color-coded.

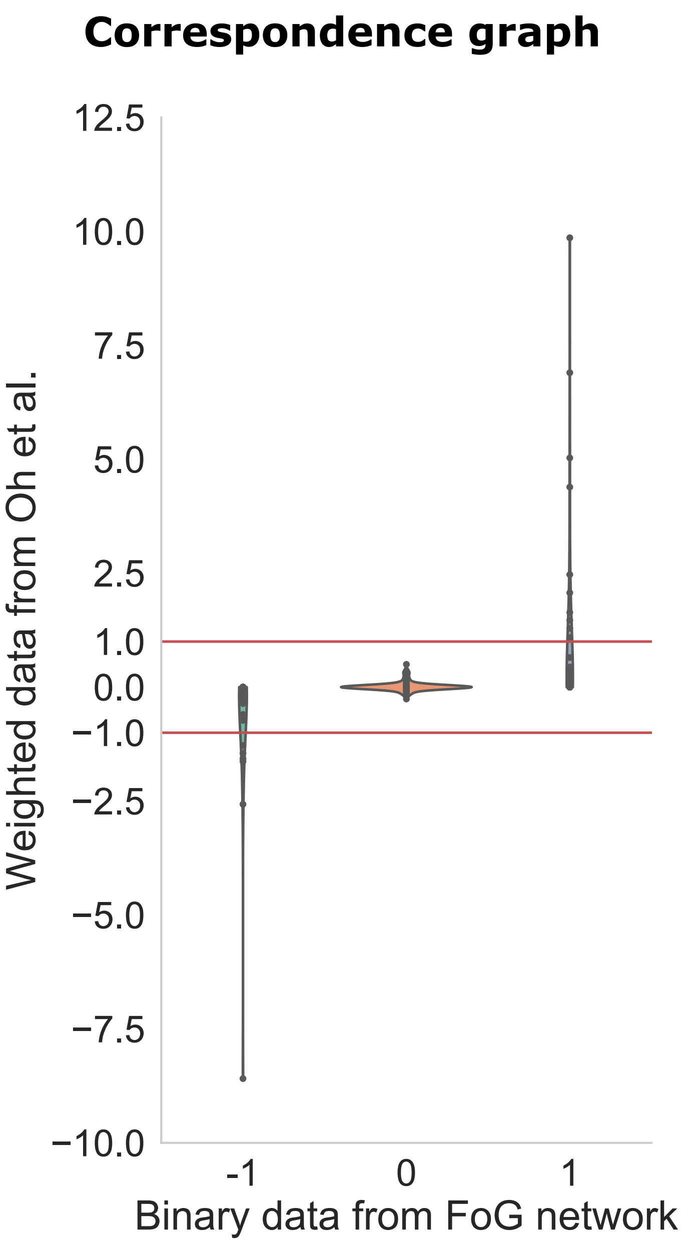


**Figure S2. Connectome comparison.**

The edge weights from (Oh et al., 2014) (Y-axis) are grouped along the binary weights of the corresponding edges used in healthy FoG network configuration (X-axis). The red lines show 1 and $-1$ weights borders.
